# Age-related differences in the loss and recovery of serial sarcomere number following disuse atrophy in rats

**DOI:** 10.1101/2024.06.10.598222

**Authors:** Avery Hinks, Geoffrey A. Power

## Abstract

**Background:** Older adults exhibit a slower recovery of muscle mass following disuse atrophy than young adults. At a smaller scale, muscle fibre cross-sectional area (i.e., sarcomeres in parallel) exhibits this same pattern. Less is known, however, about age-related differences in the recovery of muscle fibre length, driven by increases in serial sarcomere number (SSN), following disuse. The purpose of this study was to investigate age-related differences in SSN adaptations and muscle mechanical function during and following muscle immobilization. We hypothesized that old rats would experience a similar magnitude of SSN loss during immobilization, however, take longer to recover SSN than young following cast removal, which would limit the recovery of muscle mechanical function.

**Methods:** We casted the plantar flexors of young (8 months) and old (32 months) male rats in a shortened position for 2 weeks, and assessed recovery during 4 weeks of voluntary ambulation. Following sacrifice, legs were fixed in formalin for measurement of soleus wet weight and SSN with the un-casted soleus acting as a control. Ultrasonographic measurements of pennation angle (PA) and muscle thickness (MT) were also conducted weekly. *In-vivo* active and passive torque-angle relationships were constructed pre-cast, post-cast, and following 4 weeks of recovery.

**Results:** From pre- to post-cast, young and old rats experienced similar decreases in SSN (–20%, *P*<0.001), muscle wet weight (–25%, *P*<0.001), MT (–30%), PA (–15%, *P*<0.001), and maximum isometric torque (–40%, *P*<0.001), but there was a greater increase in passive torque in old (+180%, *P*<0.001) compared to young rats (+68%, *P*=0.006). Following cast removal, young exhibited quicker recovery of SSN, PA, and MT than old, but SSN recovered sooner than PA and MT in both young and old. Muscle wet weight recovered 90% and active torque fully recovered in young rats, whereas in old these remained unrecovered at 75% and 72%, respectively.

**Conclusions:** This study showed that old rats retain a better ability to recover longitudinal compared to parallel muscle morphology following cast removal, making SSN a highly adaptable, appealing mechanism for restoration of functional capacity following disuse in elderly populations.

## Introduction

Natural adult aging is associated with a decline in muscle mass, occurring at a rate of at least 0.5% per year in humans after age 60 (1–5). This loss of muscle mass is accounted for not only by losses of muscle fibre number and cross-sectional area (2,5), but also by a reduction in muscle fascicle length (FL) (6–8), which studies on animals have shown is driven by the loss of sarcomeres aligned in series (9–11). A muscle’s serial sarcomere number (SSN) is tightly coupled with its mechanical performance including force production throughout the range of motion and resting passive tension (12–14). Aging is accompanied by a reduced capacity to generate active force and an increase in resting passive tension, and the loss of SSN contributes to those mechanical impairments (5,11,15–17). Loss of SSN and strength, and increased resting passive tension are also observed following periods of immobilization with the muscle casted in a shortened position (18–21). These negative alterations to muscle morphology and function are important to consider as older adults are prone to falls and illnesses, making periods of cast-immobilization or bed rest common, and the negative effects potentially additive on top of aging (22,23), worsening their trajectory towards loss of independence.

To mitigate or reverse the loss of SSN in both aging and immobilization, interventions promoting an increase in SSN have been proposed. Studies on young adult rodents have investigated the ability for serial sarcomerogenesis through a period of simply voluntary ambulation following immobilization-induced SSN loss, and SSN recovered within 3-4 weeks following cast removal (24–26) likely due to the considerable stretch stimulus that ambulation imposed on the muscle after being immobilized in a shortened position. This stimulus for serial sarcomerogenesis may, however, be limited in old rats because a stiffer ECM limits their range of motion and fascicle stretch during walking (27). Furthermore, a decline in habitual physical activity likely contributes to the age-related loss of functional capacity in humans (28–30). Similarly, old rats exhibit lower voluntary physical activity than young rats (31), which could further limit the stimulus for re-growth during voluntary ambulation following disuse atrophy.

Previous studies have provided important insight into age-related differences in recovery following disuse atrophy. In humans, Suetta et al. (32) observed a smaller loss of quadriceps muscle volume in old compared to young men following 2 weeks of immobilization, however, after a subsequent 4 weeks of retraining, muscle volume recovered less in old than young men. In the same cohort, Suetta et al. (33) observed no recovery of vastus lateralis single fibre cross-sectional area in old men following 4 weeks of retraining, but full recovery in young men. A similar blunting of muscle mass recovery following 4 weeks of immobilization was observed in old rats (34). While these previous studies provided understandings of age-related differences in the recovery of whole-muscle mass and parallel muscle morphology (i.e., cross-sectional area) following immobilization, it is unclear whether there is also a blunted ability to recover SSN in old age, which is the driving mechanism of longitudinal muscle growth.

Comparing young (8 months) and old (32 months) Fisher 344/Brown Norway rats, the present study aimed to investigate: 1) age-related differences in the loss of soleus SSN and plantar flexor mechanical function during 2 weeks of casting in a shortened position; and 2) age-related differences in the recovery of soleus SSN and mechanical function during 4 weeks of voluntary ambulation following cast removal. We hypothesized that old rats would exhibit a smaller magnitude of SSN loss than young following immobilization, however, would take longer to recover their SSN, contributing to impaired mechanical function and blunted longitudinal muscle growth.

## Methods

### Animals

10 young (8 months) and 11 old (32 months) male Fisher 344/Brown Norway F1 rats were obtained (Charles River Laboratories, Senneville, QC, Canada). All protocols were approved by the University of Guelph’s Animal Care Committee (AUP #4905) and followed guidelines from the Canadian Council on Animal Care. Rats were housed at 23°C in groups of two or three and given ad-libitum access to a Teklad global 18% protein rodent diet (Envigo, Huntington, Cambs., UK) and room-temperature water. 5 old and 5 young rats were sacrificed after the 2-week casting intervention, and the remaining 6 old and 5 young rats were sacrificed after the 4-week recovery period following cast removal. Ultrasound measurements on the casted soleus were obtained at 7 time points: no more than 1 week prior to the application of casts (pre-cast), 1 week into the casting intervention (1 wk cast), on the day of cast removal (post-cast), and 1, 2, 3, and 4 weeks following cast removal (1, 2, 3, and 4 wk recovery). Mechanical testing measurements were obtained at three time points: pre-cast, post-cast, and 4 wk recovery. While it is well-recognized that old rats partake in less voluntary physical activity than young (31), to encourage voluntary ambulation in both groups following cast removal, rats were housed in a double-cage setup including a large tube that connects two cages together (Figure 1). Following sacrifice, the hindlimbs were fixed in formalin for subsequent determination of soleus SSN. In accordance with previous studies (19,21,35–37), the left leg served as the experimental leg while the right leg served as an internal control.

**Figure 1:**
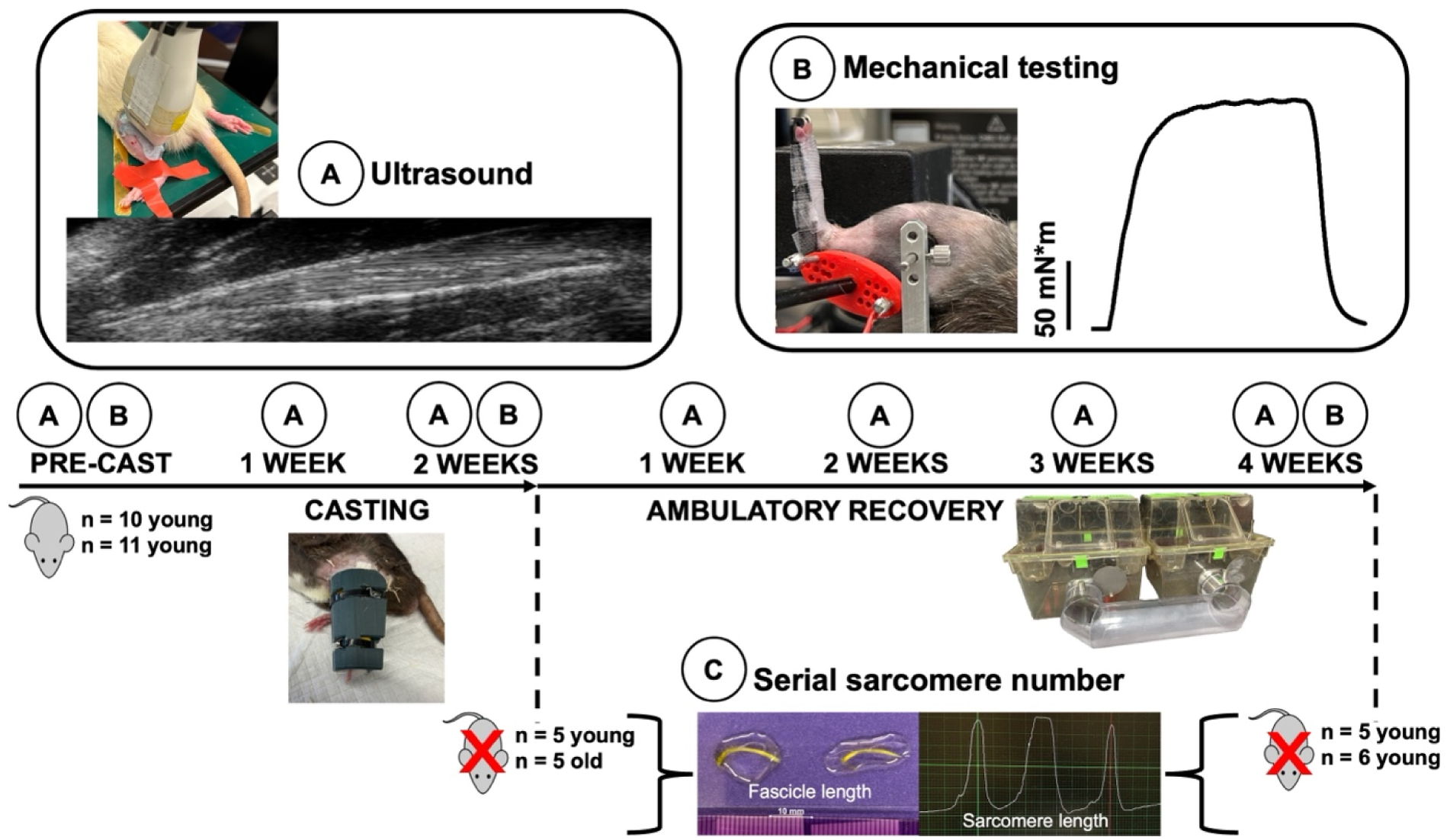
Experimental timeline. Young (n = 10) and old (n = 11) rats underwent 2 weeks of unilateral casting in full plantar flexion followed by 4 weeks of ambulatory recovery in a “double-cage” setup. Ultrasound measurements **(A)** were performed weekly. Mechanical testing **(B)** (construction of a passive and active torque-angle relationship) of the plantar flexors was performed pre-cast, after the 2-week casting intervention, and at 4 weeks of recovery. Post-cast, 5 young and 5 old rats were sacrificed for serial sarcomere number (SSN) measurements **(C)**. The remaining 5 young and 6 old rats were sacrificed after 4 weeks of recovery for SSN measurements.

### Unilateral Immobilization

Using gauze padding, vet wrap, and a 3D-printed brace and splint, the right hindlimb of each rat was casted in full plantarflexion for 2 weeks (Figure 1). The toes were left exposed to monitor for swelling (37,38). Casts were inspected daily and repaired/replaced as needed. All casts were replaced 1 week into the casting period following the 1 wk cast ultrasound measurements. Since rats were free to walk around while wearing their casts, this intervention merely promoted disuse of the immobilized muscles rather than complete unloading such as that achieved with hindlimb suspension (39).

### Data acquisition during mechanical testing and training

A 701C High-Powered, Bi-Phase Stimulator (Aurora Scientific, Aurora, ON, Canada) was used to evoke transcutaneous muscle stimulation via a custom-made electrode holder with steel galvanized pins situated transversely over the popliteal fossa and the calcaneal tendon (Figure 1). During piloting, we determined this stimulation setup produced similar values of maximum isometric tetanic torque across repeated testing sessions, and produced plantar flexion torque values consistent with values in previous literature using direct nerve stimulation (40). Torque, angle, and stimulus trigger data were sampled at 1000 Hz with a 605A Dynamic Muscle Data Acquisition and Analysis System (Aurora Scientific, Aurora, ON, Canada) with a live torque tracing during all training and mechanical data collection sessions.

### Mechanical testing

The rats were anesthetized with isoflurane and positioned on a heated platform (37°C) in a supine position. After shaving the leg completely of hair, the left leg was fixed to a force transducer/length controller foot pedal via tape, with the knee immobilized at 90°. Each mechanical testing session began with determination of the optimal current for stimulation (frequency = 100 Hz, pulse duration = 0.1 ms, train duration = 500 ms) at an ankle angle of 90° (ankle angle = angle between the tibia and the foot sole; full plantar flexion = 180°), which was the current used throughout the remainder of the session. This stimulation current was confirmed to maximally activate the plantar flexors with minimal spread to the (antagonist) dorsiflexors by completing another stimulation with the current further increased by 10 mA, in which a decrease in active torque was observed – indicating the current we used for the experiments involved minimal to no antagonist activation. 0.5-ms, 100-Hz isometric contractions were then completed at ankle angles of 70°, 80°, 90°, 100°, and 110°, each separated by 2 min of rest. Active torque was measured by subtracting the minimum value of torque at baseline (i.e., the passive torque) from the maximum value of total torque during stimulation (14,41). Following movement of the foot pedal, stimulation was preceded by 5 s of rest to reduce the impact of stress-relaxation on the measurements of passive torque.

### Ultrasonography

A UBM system (Vevo 2100; VisualSonics, Toronto, ON, Canada) operating at a centre frequency of 21 MHz was used to acquire images of the soleus, with a lateral resolution of 80 μm and an axial resolution of 40 μm (37,42). A 23-mm long probe was used, allowing acquisition of images displaying muscle fascicles from end to end. Image acquisition was optimized with an image depth of 14-15 mm for the soleus, allowing a maximum frame rate of 16 Hz (37). Prior to image acquisition, rats were anesthetized using isoflurane. With the knee fully extended, tape was used to fix the left ankle at 90° with the rat in a prone position and the hindlimb externally rotated, with the probe overlying the lateral aspect of the posterior shank (Figure 1). All ultrasound images were acquired by the same individual (A.H.). The probe position was carefully adjusted to obtain the clearest possible view of fascicles in all of the proximal, middle, and distal regions of the muscle. Throughout image acquisition, the probe was stabilized by a crane with fine-tune adjustment knobs, minimizing pressure and limiting the error associated with human movement.

Ultrasound images were analysed using ImageJ software (37,43). ImageJ’s multisegmented tool allowed careful tracing of the fascicle paths from end to end in measuring FL. Two measurements of FL and pennation angle were obtained from each of the proximal, middle, and distal regions of each muscle (i.e., n = 6 FL and pennation angle measurements per muscle) and averaged for the reporting of data, as we showed previously that soleus FL differs minimally across muscle regions (11,37). Pennation angle was defined as the angle between the fascicle and the aponeurosis at the fascicle’s distal insertion point. Three measurements of muscle thickness (proximal, middle, and distal) were also taken at the soleus mid-muscle belly (i.e., the thickest portion) (44,45). All FL, pennation angle, and muscle thickness measurements were obtained by the same experimenter (A.H.).

### Serial sarcomere number determination

Following their final mechanical testing session, rats were sacrificed via isoflurane anesthetization followed by CO_2_ asphyxiation. The hindlimbs were amputated, skinned with all muscles overlying the soleus removed, and fixed in 10% phosphate-buffered formalin with the ankle pinned at 90°. After fixation for 1-2 weeks, the soleus was dissected off the lower leg, weighed, then re-submerged in formalin until the commencement of SSN estimations. To commence the process of SSN estimations, the muscles were rinsed with phosphate-buffered saline, then digested in 30% nitric acid for 6-8 h to remove connective tissue and allow for individual muscle fascicles to be teased out (14,46).

For each muscle, two fascicles were obtained from each of the proximal, middle, and distal regions (i.e., n = 6 fascicles total per muscle). SSN and FL values were averaged across these six fascicles for the reporting of data, as we showed previously that SSN differs minimally across region in the rat soleus (11,37). Dissected fascicles were placed on glass microslides (VWR International, USA), then FLs were measured using ImageJ software (version 1.53f, National Institutes of Health, USA) from pictures captured by a level, tripod-mounted digital camera, with measurements calibrated to a ruler in plane with the fascicles (Figure 1). Sarcomere length (SL) measurements were taken at n = 6 different locations proximal to distal along each fascicle via laser diffraction (Coherent, Santa Clara, CA, USA) with a 5-mW diode laser (500 μm beam diameter, 635 nm wavelength) and custom LabVIEW program (Version 2011, National Instruments, Austin, TX, USA) (47) (Figure 1), for a total of n = 36 SL measurements per muscle. For each fascicle, the six SL measurements were averaged to obtain a value of average SL. Given the laser diameter of ∼1 mm, one SL measurement itself represents an average of thousands of SLs. Our total quantity of SL and FL measurements is consistent with previous studies (14,41,46). For each fascicle, the six SL measurements were averaged to obtain a value of average SL. Serial sarcomere number of each fascicle was calculated as:

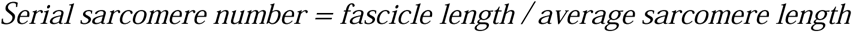

Within each fascicle, SL standard deviation (SL SD) was also noted as an estimate of SL non-uniformity.

### Estimation of time-course SSN adaptations using ultrasound-derived FL

It is often difficult to measure the time course of SSN adaptations due to the invasive nature of SSN measurements, requiring separate groups of animals to be sacrificed at each time point. We recently verified a method of using ultrasonographic measurements of FL to estimate SSN adaptations in the rat soleus provided that a correction factor is applied (37). Thus, as a final analysis to address our primary research question (age-related differences in SSN adaptations during and following immobilization), we used the ultrasound-derived measurements of FL to estimate the time course of SSN adaptations. To do this, we first calculated the ratio of the ultrasound-derived FL values to the measures of FL on dissected fascicles (this value was 1.04; see Results). This ratio was used as a correction factor to estimate what dissected FL would be at the 1 wk cast and 1, 2, and 3 wk recovery time points. The control leg measurements of SSN were used as estimates of SSN pre-cast and the SSN measurements at 4 wk recovery represented 4 wk recovery. At post-cast, only half the rats (the n = 5 young and n = 5 old that were sacrificed at that time point) had SSN measurements performed, so the remaining rats had SSN estimated at that time point using the ultrasound-derived FL correction factor. Since SL did not differ between the post-cast and 4 wk recovery time points in old or young rats (see Results), SL values from the experimental leg were used for estimations of SSN post-cast and at 1, 2, and 3 wk recovery. To estimate SL at 1 wk cast, we took the average of the control and experimental leg SL and used that in estimation of SSN. To summarize, estimations of SSN at time points for which we did not perform SSN measurements were done via the equation:

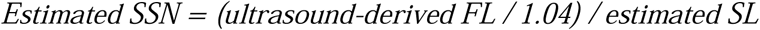

### Statistical analysis

All statistical analyses were performed in SPSS Statistics Premium 28. To investigate baseline age-related differences in muscle structure, one-way analysis of variance (ANOVA) was used on pre-cast measurements for ultrasound data (ultrasound-derived FL, pennation angle, and muscle thickness) and on measurements from the control leg for dissected muscle data (muscle wet weight, SSN, SL, SL SD, dissected FL). To investigate differences between time points in each age group for dissected muscle measurements, two-way repeated measures ANOVA (leg [control, casted] × time [post-cast, 4 weeks recovery]) were employed. Subsequently, to investigate age-related differences in adaptations to casting and recovery for the dissected muscle measurements, two-way ANOVAs (age [young, old] × time [post-cast, 4 weeks recovery]) were employed on the data normalized to control leg values. To investigate age-related differences in the time course of ultrasound measurements, two-way repeated measures ANOVA (age [young, old] × time [pre-cast, 1 week cast, post-cast, 1, 2, 3, 4 weeks recovery]) were employed on the data normalized to pre-cast. To investigate changes in the active and passive torque-angle relationships, three-way repeated measures ANOVA (age [young, old] × time [pre-cast, post-cast, 4 weeks recovery] × angle [70°, 80°, 90°, 100°, 110°]) were employed. Lastly, to evaluate the estimated time-course changes in SSN (as estimated via ultrasound-derived FL) throughout the whole study, we applied a two way ANOVA (age [young, old] × time point [pre-cast, 1 wk cast, post-cast, 1, 2, 3, 4 wk recovery]).

Where main effects or interactions were detected, two-tailed t-tests were used for pairwise comparisons, with a Sidak correction for multiplicity. Significance was set at α = 0.05. All values are reported as the mean ± standard deviation.

## Results

### Baseline age-related differences in soleus muscle morphology

Muscle wet weight, SSN, and dissected FL as measured at 90° were 8%, 11%, and 12% less, respectively, in the control legs of old compared to young rats (Table 1). SL and SL SD as measured at 90° did not differ between young and old rats (Table 1). From the ultrasound measurements at 90° pre-cast, FL and muscle thickness were 8% and 15% less, respectively in old compared to young rats, while pennation angle did not differ between old and young (Table 1).

**Table 1:**
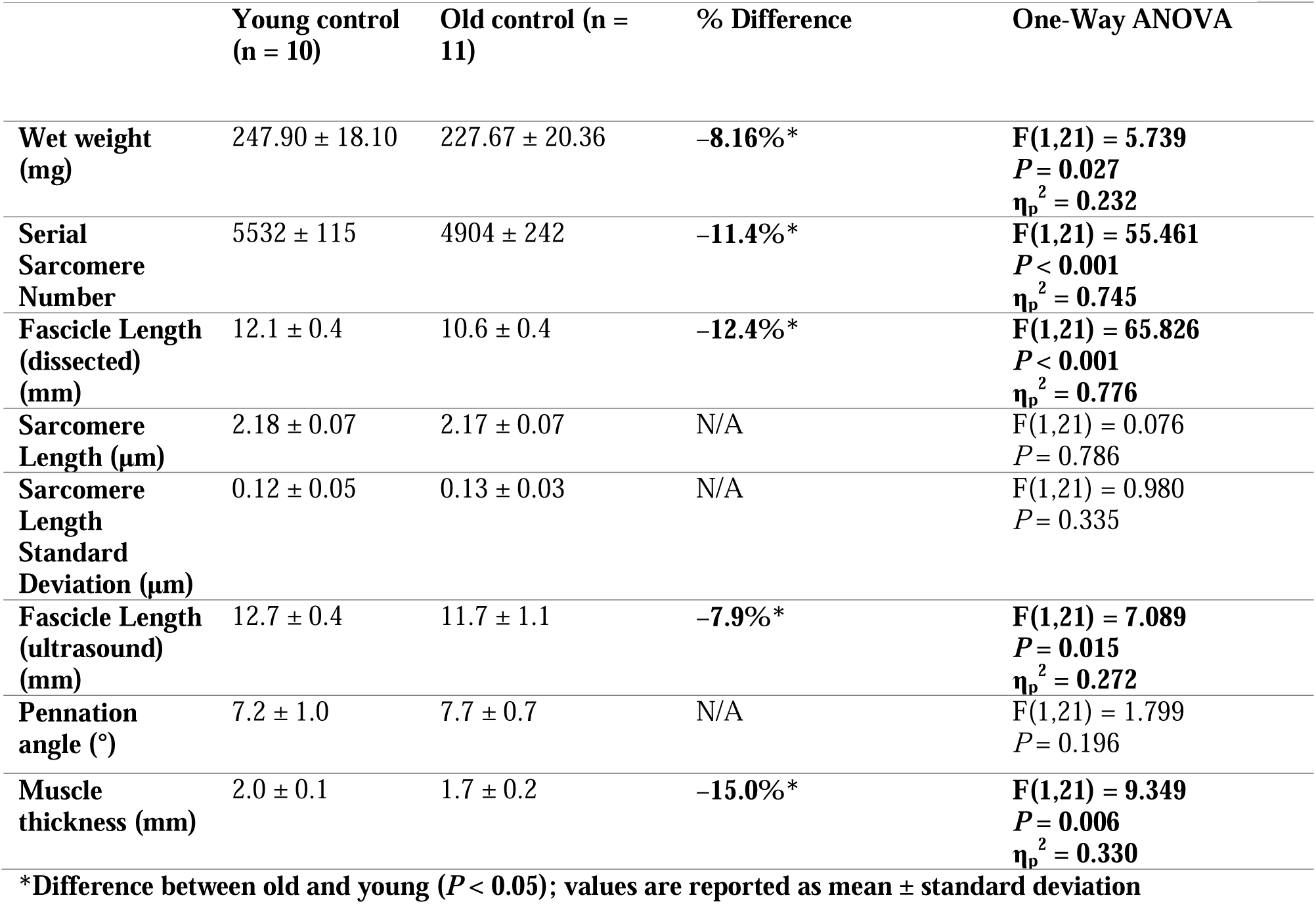
Baseline soleus morphological properties in old compared to young rats.

### Influence of immobilization and recovery on muscle wet weight in young and old rats

For muscle wet weight of young rats, there was a leg × time interaction (F(1,8) = 12.121, *P* = 0.008, η_p_^2^ = 0.602). Post-cast, young rats had a 27% lower muscle wet weight in the casted compared to control leg (*P* < 0.001) (Figure 2A). At 4 wk recovery, muscle wet weight of young rats was then greater in the casted leg compared to post-cast (*P* < 0.001), however, was still less than that of the control leg by 10% (*P* = 0.008) (Figure 2A). By contrast, old rats only showed an effect of leg (F(1,9) = 55.621, *P* < 0.001, η_p_^2^ = 0.861) for muscle wet weight such that muscle wet weight of the casted leg was 25% less than that of the control leg both post-cast and at 4 wk recovery (Figure 2A). Indeed, when analyzing the casted leg values as a percent of control leg values, there was an age × time interaction (F(1,21) = 5.797, *P* = 0.028, η_p_^2^ = 0.254): old and young showed no difference (*P* = 0.641) in the initial reduction to 72-75% of control values post-cast, but young recovered to 90% of control values at 4 wk recovery (*P* = 0.004 compared to post-cast) while old remained at 75% and did not differ from post-cast (*P* = 0.967) (Figure 2B).

**Figure 2:**
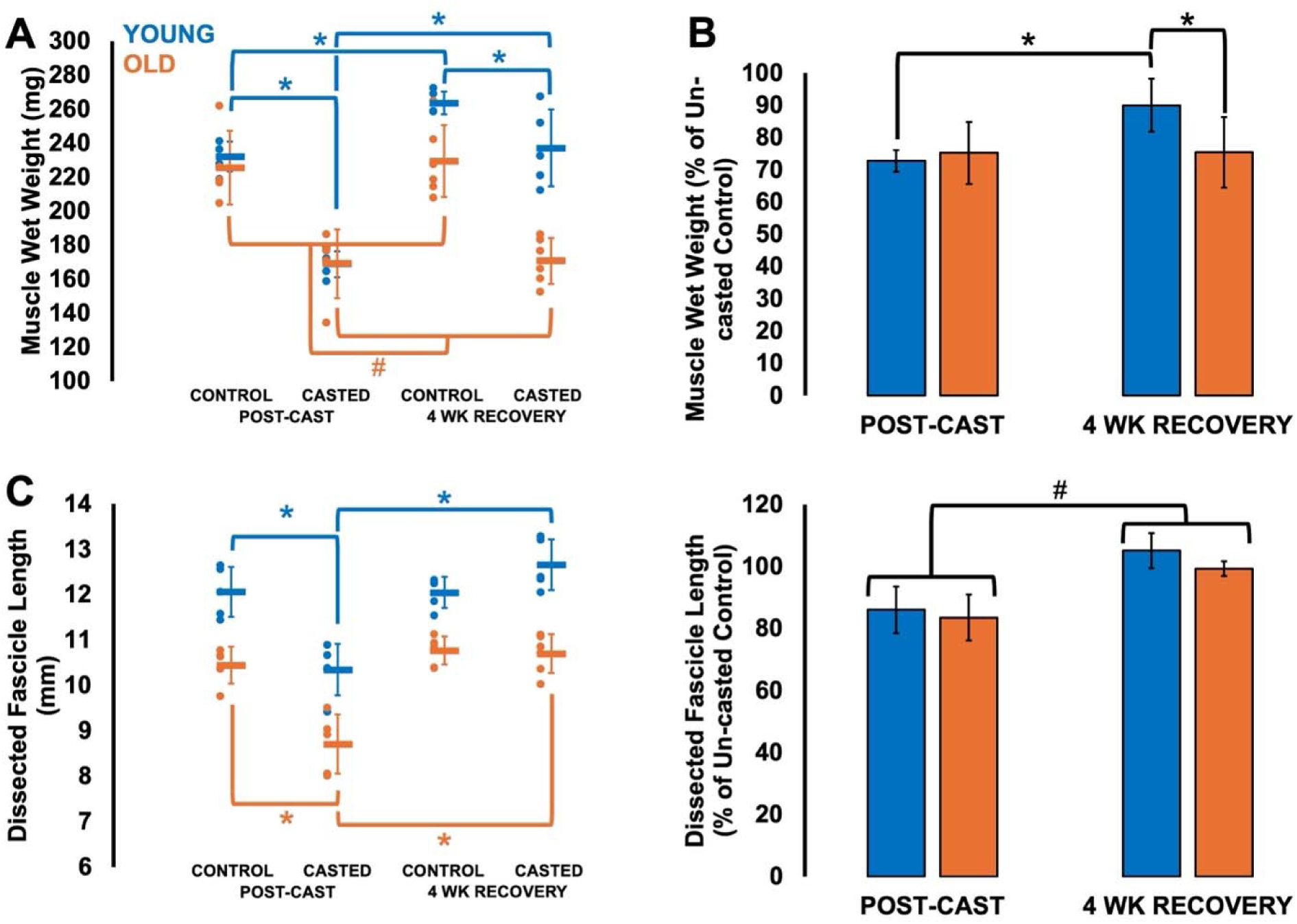
Differences in muscle wet weight (**A-B**) and fascicle length (**C-D**) of dissected fascicles as measured at 90° at post-cast (n = 5 young, n = 5 old) and 4 weeks of recovery (n = 5 young, n = 6 old) compared to control muscles shown as absolute values (**A and C**) and with experimental muscles expressed as a percent of control muscles (**B and D**). Data are displayed as mean ± standard deviation. *Difference between indicated points (*P* < 0.05). #Effect of leg (A) or age (C) with points combined (*P* < 0.05).

### Influence of immobilization and recovery on fascicle length and sarcomere length as measured at 90° from dissected muscle in young and old rats

For dissected FL as measured at 90°, there were leg × time interactions for both young (F(1,8) = 18.633, *P* = 0.003, η_p_^2^ = 0.700) and old (F(1,8) = 23.369, *P* < 0.001, η_p_^2^ = 0.722), which showed similar decreases in FL from the control to the casted leg post-cast (young: -14%, *P* = 0.002; old: -17%, *P* < 0.001) (Figure 2C). FL of the casted leg then increased from post-cast to 4 wk recovery in young (*P* < 0.001) and old (*P* < 0.001) and did not differ from the control leg (young: *P* = 0.148; old: *P* = 0.766) (Figure 2C). When analyzing the casted leg values as a percent of pre-cast, there was an effect of time (F(1,21) = 44.951, *P* < 0.001, η_p_^2^ = 0.726) but not age (F(1,21) = 2.550, *P* = 0.129), suggesting similar recovery of FL by 4 weeks in both groups (Figure 2D).

In young and old rats, SL as measured at 90° showed no effects of time (young: F(1,9) = 0.047, P = 0.834; old: F(1,9) = 0.570, *P* = 0.470) but did show effects of leg (young: F(1,8) = 23.888, *P* = 0.001, η_p_^2^ = 0.749; old: F(1,9) = 7.575, *P* = 0.022, v = 0.457), with SL being longer in the casted leg compared to the control leg in young (control: 2.18 ± 0.07 μm; casted: 2.38 ± 0.12 μm) and old rats (control: 2.17 ± 0.07 μm; casted: 2.25 ± 0.08 μm) regardless of time point (Supplemental Figure S1A). The same pattern was observed for SL SD, with no effects of time (young: F(1,8) = 0.123, *P* = 0.735; old: F(1,9) = 3.402, *P* = 0.098) but effects of leg (young: F(1,8) = 8.168, *P* = 0.021, η_p_^2^ = 0.505; old: F(1,9) = 9.941, *P* = 0.012, η_p_^2^ = 0.525) that showed greater SL SD (young: +59%; old: +32%) in the casted than the control leg regardless of time point (Supplemental Figure S1B). Interestingly, SL SD explained 34% of the variation in SL across all control and casted muscles (*P* < 0.001) (Supplemental Figure S1C), but when separated out into each leg and age group, SL SD explained 81% of the variation in SL in the casted leg of young rats (*P* < 0.001), but these two variables did not relate in control legs of young rats (*P* = 0.860) nor control (*P* = 0.885) or casted legs (*P* = 0.653) of old rats (Supplemental Figure S1D-G). Therefore, these longer SLs seemed to be mostly accounted for by nonuniformity in SL at rest, especially for casted muscles from young rats.

### Influence of immobilization and recovery on ultrasound-derived fascicle length, pennation angle, and muscle thickness in young and old rats

For ultrasound-derived FL, there was an age × time interaction (F(6,54) = 5.357, *P* < 0.001, η_p_^2^ = 0.373). Old rats showed a greater reduction in FL at 1 wk cast compared to young (*P* = 0.030), but the magnitude of FL decrease post-cast did not differ between young and old rats (*P* = 0.593) despite young decreasing by 15% (*P* < 0.001) and old decreasing by 22% (*P* < 0.001) (Figure 3B). Ultrasound-derived FL also recovered sooner in young rats, as they no longer differed from pre-cast at 1 wk recovery (*P* = 1.000), while old rats differed from pre-cast at 1 wk (*P* = 0.005) and 2 wk (*P* = 0.013), then recovered at 3 wk (*P* = 0.075 compared to pre-cast) (Figure 3B). Correspondingly, ultrasound-derived FL relative to pre-cast was also ∼12% less in old compared to young rats at 1, 2, 3, and 4 wk recovery (*P* = 0.006-0.022) (Figure 3B).

**Figure 3:**
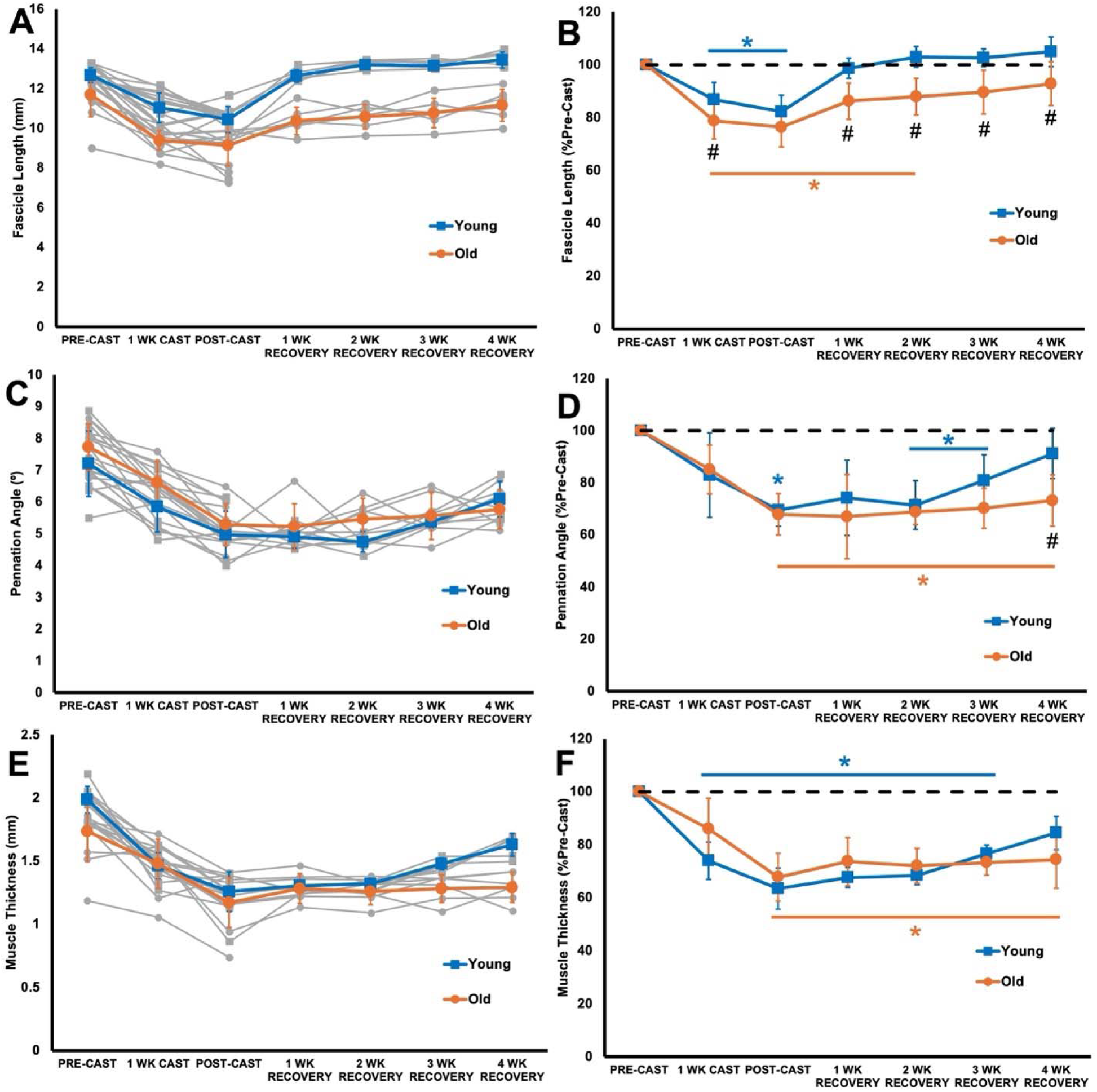
Changes in ultrasound-derived fascicle length (**A-B**), pennation angle (**C-D**), and muscle thickness (**E-F**) throughout casting (n = 10 young, n = 11 old) and recovery (n = 5 young, n = 6 old), with half the rats sacrificed post-cast and the remaining half sacrificed after 4 weeks of recovery. Left graphs display absolute values with individual data (grey lines). Right graphs show data normalized to pre-cast, on which statistical analyses were performed. Data are displayed as mean ± standard deviation. *Difference from pre-cast (*P* < 0.05). #Difference between young and old (*P* < 0.05).

For pennation angle, there was also an age × time interaction (F(6,54) = 2.540, *P* = 0.031, η_p_^2^ = 0.220). Pennation angle of young (*P* = 0.466) and old rats (*P* = 0.275) did not significantly differ from pre-cast at 1 wk cast, but decreased by post-cast (young: -17%; old: -13%; both *P* < 0.001) (Figure 3D). The magnitude of decrease did not differ between old and young (*P* = 0.827). Old rats also did not differ from young in pennation angle relative to pre-cast at 1, 2, and 3 wk recovery (*P* = 0.072-0.458), but at 4 wk recovery, young rats were greater than old by 18% (*P* = 0.014) (Figure 3D). Correspondingly, pennation angle of young rats was unrecovered at 2 (*P* < 0.001) and 3 wk (*P* = 0.017) then recovered by 4 wk recovery (*P* = 0.810 compared to pre-cast), but in old rats remained reduced compared to baseline across the entire recovery period (*P* < 0.001-0.011 compared to pre-cast) (Figure 3D).

For muscle thickness, there was also an age × time interaction (F(6,54) = 4.958, *P* < 0.001, η_p_^2^ = 0.355), however, muscle thickness relative to pre-cast did not differ between young and old at any time points (*P* = 0.068-0.337) (Figure 3F). At 1 wk cast, muscle thickness decreased in young rats (-23%, *P* = 0.018) but not old rats (*P* = 0.652). At post-cast, young decreased further (-37%, *P* < 0.001) and old decreased compared to pre-cast (-30%, *P* < 0.001) (Figure 3F). Muscle thickness of young rats differed from pre-cast at 1, 2, and 3 wk recovery (all *P* < 0.001) then no longer differed from pre-cast at 4 wk (*P* = 0.079), while muscle thickness of old rats differed from pre-cast throughout the entire recovery period (*P* < 0.001-0.001) (Figure 3F).

### Adaptations in serial sarcomere number during casting and recovery in young and old rats

For SSN, there was a leg × time interaction for both young (F(1,8) = 130.651, *P* < 0.001, η_p_^2^ = 0.942) and old (F(1,9) = 28.777, *P* < 0.001, η_p_^2^ = 0.762). Young (-22%, *P* < 0.001) and old (-19%, *P* < 0.001) showed similar reductions in SSN from the control to casted leg post-cast (Figure 4A). SSN of the casted leg in both young (*P* < 0.001) and old (*P* < 0.001) then increased from post-cast to 4 wk recovery, however, in young the casted and control legs no longer differed in SSN (*P* = 0.084), while SSN of the casted leg in old rats was still 4% less than the control leg (*P* = 0.046) (Figure 4A). With that said, when analyzing the casted leg values as a percent of control leg values, there was only an effect of time (F(1,21) = 121.861, *P* < 0.001, η_p_^2^ = 0.878) and no effect of age (F(1,21) = 0.081, *P* = 0.779), which would suggest SSN recovered similarly by 4 weeks in young and old rats (Figure 4B).

**Figure 4:**
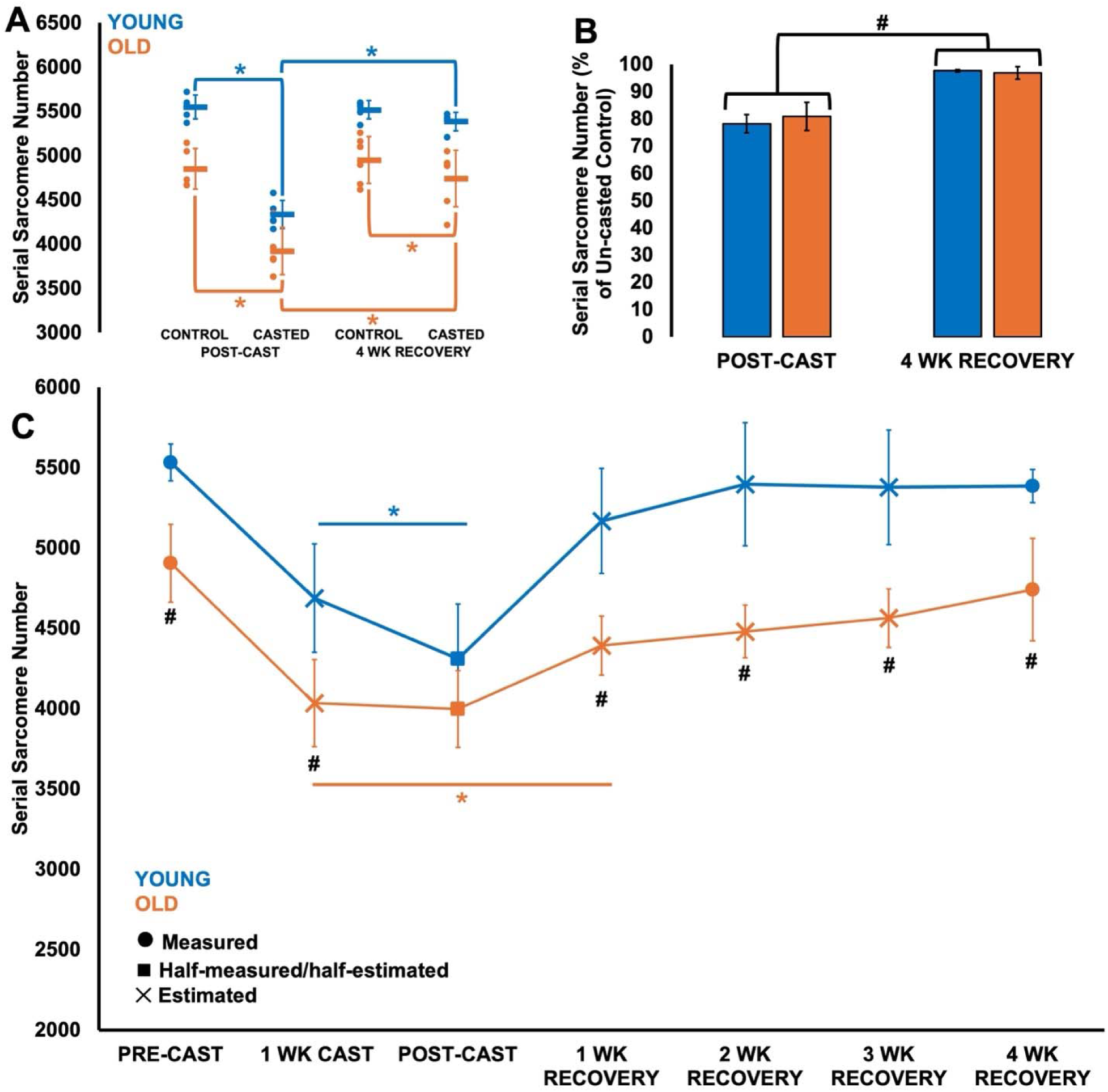
**A-B.** Differences in serial sarcomere number at post-cast (n = 5 young, n = 5 old) and 4 weeks of recovery (n = 5 young, n = 6 old) compared to control muscles, shown as absolute values (A) and with experimental muscles expressed as a percent of control muscles (B). Data are displayed as mean ± standard deviation. *Significant difference between indicated points (*P* < 0.05). #Effect of leg with points combined (*P* < 0.05). **C.** Time-course changes in serial sarcomere number throughout casting (n = 10 young, n = 11 old) and recovery (n = 5 young, n = 6 old), estimated using ultrasound-derived fascicle length. Data are displayed as mean ± standard deviation. *Difference from pre-cast (*P* < 0.05). #Difference between young and old (*P* < 0.05).

Ultrasound-derived FL was on average 1.04 times FL of dissected fascicles from the same muscles (Supplemental Figure S2). When using ultrasound-derived FL to estimate time-course changes in SSN, there was an age × time interaction (F(6,54) = 2.739, *P* = 0.021, η_p_^2^ = 0.233). Old had a lower SSN than young at all time points (*P* <0.001-0.002) expect for post-cast (*P* = 0.339), suggesting sarcomere loss reached its limit earlier in old (1 week into casting) than in young rats (2 weeks into casting) (Figure 4C). Young only had a lower SSN compared to pre-cast at 1 wk cast and post-cast (both *P* < 0.001), no longer differing from pre-cast at 1 wk recovery and onwards (*P* = 0.450-1.000) (Figure 4C). By contrast, old had a lower SSN compared to pre-cast at 1 wk cast (*P* < 0.001), post-cast (*P* = 0.003), and at 1 wk recovery (*P* = 0.027), and no longer differed from pre-cast at 2 wk onwards (*P* = 0.054-0.373) (Figure 4C). These data suggest SSN of young rats recovered earlier (1 week following cast removal) than old rats (2 weeks following cast removal).

### Influence of immobilization and recovery on active and passive torque-angle relationships in young and old rats

For the active torque-angle relationship, there was an age × time × angle interaction (F(8,72) = 7.487, *P* < 0.001, η_p_^2^ = 0.355). Old rats produced lower active torque than young at all angles pre-cast (-31 to -38%, all *P* = < 0.001), post-cast (-39 to -41%, all *P* < 0.001), and at 4 wk recovery (-52 to -56%, all *P* < 0.001) (Figure 5A). In young rats, active torque decreased at all angles from pre to post-cast (-31 to -39%, all *P* < 0.001), notably to values that were almost identical to those of old rats pre-cast (Figure 5A). Active torque then increased at all angles from post-cast to 4 wk recovery (+56-64%, all *P* < 0.001), no longer differing from pre-cast (*P* = 0.090-0.980). At pre-cast and 4 wk recovery, active torque differed among all angles (all *P* < 0.001), while at post-cast active torque was statistically the same between some angles (*P* = 0.052-0.969) (Figure 5A), suggesting a shift of the torque-angle relationship’s plateau region to a more plantar flexed angle from pre to post-cast, then a return to a more dorsiflexed angle at 4 wk recovery.

**Figure 5:**
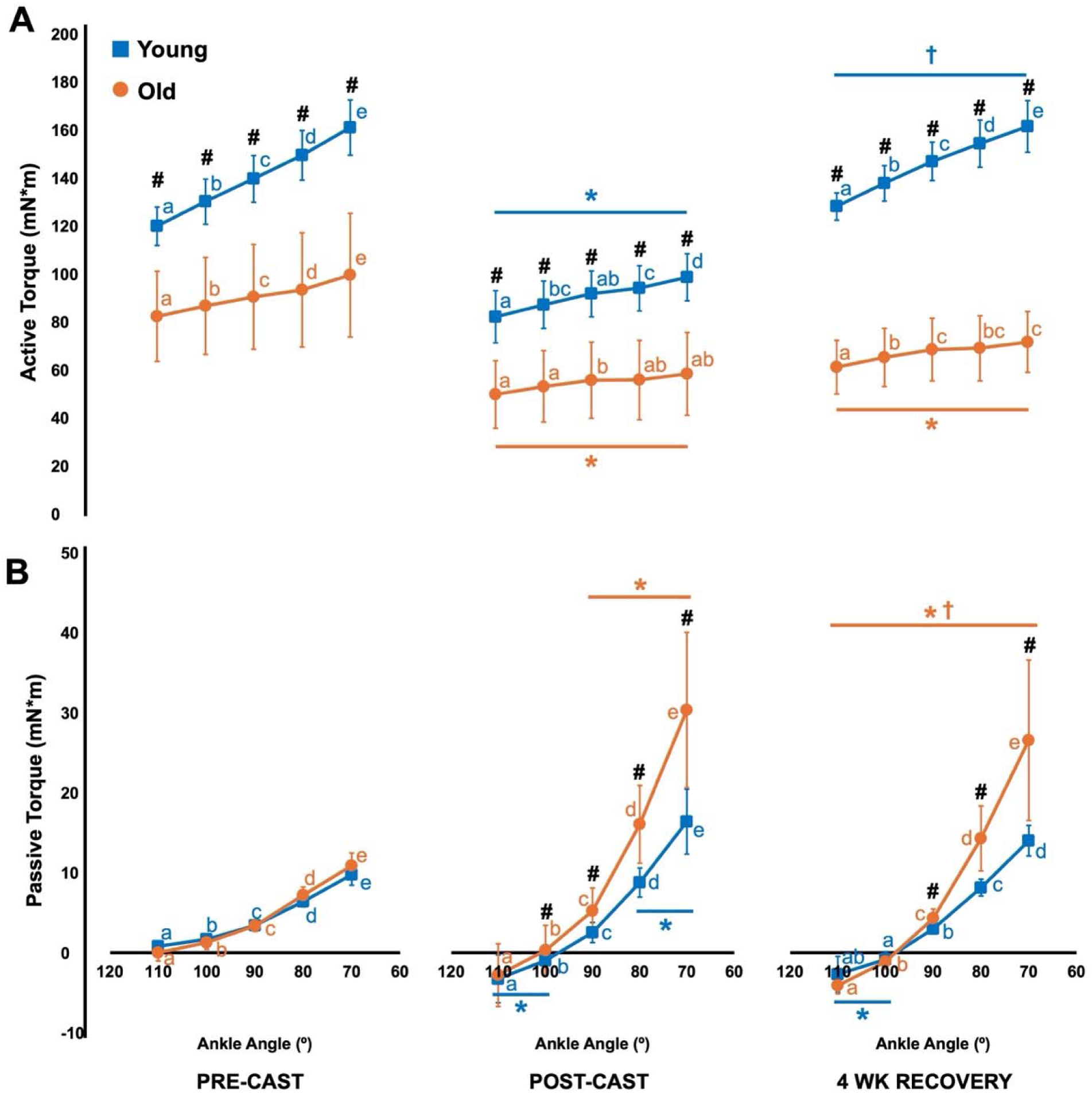
Differences in the active (**A**) and passive (**B**) torque-angle relationships between young (n = 10 pre- and post-cast; n = 5 at 4 wk recovery) and old rats (n = 11 pre- and post-cast; n = 6 at 4 wk recovery) and across time. *Difference from pre-cast (*P* < 0.05). †Difference from post-cast (*P* < 0.05). #Difference between young and old (*P* < 0.05). Same letters denote no significant difference within a time point and age group (*P* > 0.05).

In old rats, active torque also decreased at all angles from pre to post-cast (-39 to -41%, all *P* < 0.001) (Figure 5A). At 4 wk recovery, active torque at all angles was still less than pre-cast (-24 to -28%, all *P* < 0.001), and did not differ from post-cast (*P* = 0.114-0.590), indicating minimal recovery of active torque (Figure 5A). Furthermore, like in young rats, active torque differed among all angles pre-cast (*P* < 0.001-0.005), suggesting optimal torque occurred at a more dorsiflexed angle, but post-cast active torque was the same among most angles (*P* = 0.052-0.995), suggesting a shift of the plateau region to a more plantar flexed angle (Figure 5A). At 4 wk recovery, there seemed to be some reversal of this plateau region shift, with torque at 70° being greater than torque at 110° (*P* = 0.008) and 100° (*P* = 0.017) (Figure 5A).

For the passive torque-angle relationship, there was an age × time × angle interaction (F(8,72) = 4.703, *P* < 0.001, η_p_^2^ = 0.343). In both old and young rats at all time points, passive torque consistently increased as the ankle angle became more dorsiflexed (*P* < 0.001-0.024) (Figure 5B). Pre-cast, old and young rats did not differ in passive torque at any angles (*P* = 0.086-0.623). From pre- to post-cast, passive torque of young rats increased at 70° (+68%, *P* = 0.006) and 80° (+37%, *P* = 0.022), and of old rats increased at 70° (+180%), 80° (+124%), and 90° (+55%) (all *P* < 0.001) (Figure 5B). This casting-related increase in passive torque was more pronounced in old rats, as post-cast, old rats produced greater passive torque than young at angles 70° to 90° (+83-109%, P < 0.001-0.015) (Figure 5B). At 4 wk recovery, passive torque of young rats no longer differed from pre-cast at 70° (*P* = 0.482) and 80° (*P* = 0.452), however, also did not differ from post-cast (*P* = 0.165-0.448), suggesting incomplete recovery (Figure 5B). Passive torque of old rats at 4 wk recovery decreased compared to post-cast at 70° (*P* = 0.016), 80° (*P* = 0.023), and 90° (*P* = 0.019), but these were still elevated compared to pre-cast (+27- 144%, *P* < 0.001-0.007) (Figure 5B). Furthermore, old rats still produced greater passive torque than young at angles 70° to 90° at 4 wk recovery (+43-90%, *P* = 0.010-0.049).

It is also important to note that passive torque in young rats became more negative at 100° and 110° post-cast and at 4 wk recovery (*P* < 0.001-0.012), and in old rats at 4 wk recovery (P < 0.001-0.001) (Figure 5B), suggesting a steepening of the dorsiflexor passive torque-angle curve as well.

## Discussion

The purpose of this study was to investigate age-related differences in muscular adaptations during casting in a shortened position and subsequent recovery, with a particular focus on the regulation of longitudinal muscle growth governed by SSN. We found similar magnitudes of decrease between young and old rats for SSN, muscle wet weight, muscle thickness, pennation angle, and isometric active torque production after 2 weeks of casting, but there was a greater increase in passive torque in old rats compared to young rats. Following cast removal, young rats exhibited quicker recovery of SSN, pennation angle, muscle thickness, muscle wet weight, and maximum isometric torque than old. Additionally, voluntary ambulation provided a stronger stimulus for SSN growth than growth of parallel muscle morphology (pennation angle, muscle thickness) in both young and old rats, highlighting the urgency that the system places on regulating muscle length through the addition of SSN.

### Age-related differences in baseline muscle morphology

The control soleus of old rats had a lower muscle wet weight (–8%) and SSN (–11%) compared to young. Since SL as measured at an ankle angle of 90° did not differ between young and old, the lower SSN in old rats reflected the shorter FL of dissected fascicles (–11%) as measured at 90°. Ultrasonographic measurements at the pre-cast timepoint showed similar age-related differences, with muscle thickness (–15%) and FL (–8%) both being smaller in old rats. These age-related changes in muscle morphology are consistent with previous reports in F344/BN rats (10,40,48,49), and collectively reflect an age-related loss of muscle contractile tissue, which is driven by largely motor neuron loss (50) and increased rates of protein degradation and apoptosis (48,51).

Notably, ultrasound-derived FL measurements underestimated the difference in soleus SSN between young and old rats by ∼3%. This finding aligns with our previous study that showed ultrasound-derived FL alone (i.e., without a measurement of SL), while often used as an *in-vivo* proxy of SSN, does not perfectly reflect actual SSN adaptations due to limitations associated with assuming SL, intramuscular connective tissue, and the two-dimensional nature of ultrasound scans (37). This disconnect between ultrasound-derived FL and actual SSN explains why some studies in humans have observed age-related differences in ultrasound-derived FL (6,16,53–62) while others observed no differences (63–69). Our results are consistent with previous studies that observed 7-37% lesser SSN in old than young rats and mice (9,11,70).

### Age-related differences in muscular adaptations to casting

Several studies on humans and rodents have shown that older individuals experience similar or smaller magnitude losses of muscle mass, cross-sectional area, and strength compared to young following immobilization (32–34,48,71–74). Aligning with those studies, we observed similar magnitude reductions in muscle wet weight, muscle thickness, and maximum plantar flexor torque between old and young rats following 2 weeks of immobilization in a shortened position.

Across studies on the regulation of SSN, it is generally accepted that subtraction of sarcomeres during casting in a shortened position occurs to reduce sarcomeric compression and restore the original resting SL for optimal force production in that shortened position (13,18–21). We used ultrasound-derived FL to better elucidate the time-course of immobilization-induced SSN loss, and showed that SSN loss plateaued at 1 week of immobilization in old rats, but in young rats decreased further from 1 to 2 weeks of immobilization. These findings align with Fisher & Brown (72)’s conclusion in comparing immobilization-induced loss of muscle mass between old and very old rats: immobilization had less of an effect when superimposed on top of the muscle mass loss that already occurs due to aging. In other words, young rats may simply have more sarcomeres to lose before they reach the limit of total possible SSN loss, while old reach that limit earlier due to having already lost SSN with age.

The loss of contractile tissue in parallel indicated by the reductions in muscle mass (– 25%), muscle thickness (–30% to –37%) and pennation angle (–13% to –17%) corresponded to the reductions in maximum isometric strength (–30% to –40%) in both young and old rats. Additionally, the change in shape of the active torque-angle relationship from pre- to post-cast reflected the ∼20% loss of SSN. Pre-cast, the ankle angles we assessed (70° to 110°) represented the ascending limb of the torque-angle relationship as evidenced by the significant differences in torque between each angle (Figure 5A first panel). Post-cast, however, across the same range of joint angles, we were testing more on the torque-angle relationship’s plateau region as there was more homogeneity in torque between angles (i.e., a flatter torque-angle relationship; Figure 5A second panel). This change represents a shift in optimal angle to a more shortened position, which is consistently observed alongside a decrease in SSN to optimize myofilament overlap and force production at the new resting muscle length (19–21).

While passive tension generated within the sarcomere (i.e., by the giant elastic protein titin) contributes to some of the passive torque exhibited at the joint level, passive tension generated by the extracellular matrix (ECM) seems to contribute more (75,76). Regardless, from pre- to post-cast, both young and old exhibited increases in passive torque in accordance with previous studies (18,19), however, the increase was more pronounced in old, with old rats producing 109% greater passive torque than young at 70° (i.e., the most stretched angle tested). This age-related greater increase in passive torque with casting largely reflects an age-related greater collagen accumulation and in particular greater collagen crosslinking in the ECM (77–81). The loss of SSN may have also contributed to the greater passive torque, as SL at 90° was 4-9% longer in the casted than the control leg at both post-cast and 4 wk recovery, which was driven by a greater variability in resting SL (Supplemental Figure S2). Longer resting SLs in the casted soleus would increase passive tension, which could elevate total plantar flexor passive torque. However, based on our passive torque data following the recovery period (discussed below) and the age-related ECM changes observed in previous studies, it is likely that ECM adaptations contributed more than SSN adaptations to these immobilization-induced changes in the passive torque-angle relationship.

### Age-related differences in muscular adaptations during recovery from casting

In contrast to the similar impairments in muscle morphology and function between young and old following 2 weeks of casting, old rats exhibited slower or incomplete recovery for every variable measured. By 4 wk recovery, muscle wet weight of young rats recovered to 90% of control values, but in old rats remained unrecovered at 75%, about the same as that observed post-cast (Figure 2B). Pennation angle and muscle thickness also recovered by 4 wk in young but not old rats. As measures of parallel muscle morphology (including muscle thickness, pennation angle, and to an extent muscle wet weight) are associated with maximal force production (82), it is understandable that maximal isometric torque of young rats fully recovered by 4 wk but for old rats remained 24-28% depressed compared to pre-cast and did not differ from post-cast. These age-related slower recoveries of muscle wet weight, muscle thickness, pennation angle, and maximal isometric torque are consistent with previous studies in rodents and humans (32–34,49).

Previous studies on young rodents have shown that SSN can recover within 3 weeks following cast removal (24–26). By using ultrasound-derived FL to estimate time-course changes in SSN, we demonstrated that SSN likely recovers even sooner, at least 1 week following cast removal—notably faster than the recovery of muscle thickness and pennation angle. The adaptability of SSN during recovery following disuse, like the other measures described above, was age dependent. SSN of old rats appeared to recover at 2 wk rather than 1 wk recovery (Figure 4C). With that said, this may not have represented full recovery for old rats, as SSN of the casted leg was still significantly 4% less than the control leg at 4 wk recovery in old rats (Figure 4A). Furthermore, when observing the trajectory of SSN throughout the recovery period in Figure 4C, while young rats plateaued after 1 wk recovery, old rats demonstrated a consistent upward slope across 1 wk to 4 wk recovery. Regardless, the recovery of longitudinal muscle morphology (whether ultrasound-derived FL or actual SSN) occurred faster than measures associated with parallel muscle morphology (muscle thickness and pennation angle) following cast removal in both young and old. From these findings, it appears that early on during remobilization the stimulus for SSN growth (stretching of muscle fascicles while walking) was stronger than the stimulus for parallel growth (loading of the muscle) (82).

Changes in the shape of the active torque-angle relationship at 4 wk recovery also reflected the observed SSN adaptations. The active torque-angle relationship of young returned to its original shape of the ascending region (Figure 5A third panel), which aligns with the restoration of SSN. Old’s active torque-angle relationship retained a flatter appearance, but with less homogeneity in active torque between angles than at post-cast (e.g., with torque at 70° differing from 100° and 110°), suggesting a partial shift back toward the original optimal angle. The passive torque-angle relationships at 4 wk recovery, by contrast, were similar to post-cast for both old and young rats. Since SSN recovered in both groups, these steeper passive torque-angle relationships at 4 wk of recovery are likely associated with a lack of reversal of ECM adaptations.

Old rats partake in less voluntary physical activity than young rats (31), and reduced activity would limit mechanical loading on the hindlimb and the associated stimuli for muscle growth. Therefore, the present study’s design did not separate the contributions of an age-related reduction in voluntary activity and aging itself on the old rats’ slower recovery following disuse. With that said, several factors in aged muscle could limit the stimuli and signalling for SSN regrowth following disuse. Most notably, Horner et al. (27) recently showed that greater ECM-associated passive forces in muscles of old rats limit muscle excursions during walking and the forces generated at the ends of the range of motion. Such a reduced excursion could have resulted in a weaker stimulus for serial sarcomere addition (83) in our old rats during the recovery period. Additionally, Fuqua et al. (49) found that translational capacity and myofibrillar protein synthesis were not impaired in old rats during recovery from hindlimb unloading, thus they attributed the inability to recover muscle mass to higher rates of protein degradation. Regarding specifically SSN, downregulation of MuRF1 and the ubiquitin-proteasome system (a regulator of protein degradation) contributes to serial sarcomerogenesis (84), and greater MuRF activity has been reported in older individuals (85), especially in more severe cases of sarcopenia (86). Old rats also exhibit elevated levels of apoptosis at baseline than young rats (48). These processes collectively would act against the synthesis of contractile proteins, slowing recovery of SSN. Satellite cells of aged muscle also exhibit dysfunction (87–89) and blunted responsiveness to mechanical stimuli (8,90) that limit the capacity for muscle regeneration, including during recovery from disuse (33). Lastly, old rats are more susceptible to muscle damage during recovery from disuse as compared to young rats (91), and this greater incurrence of muscle damage may result in a longer recovery period to fully restore contractile tissue (11,92).

## Conclusion

Here we showed for the first time that longitudinal muscle morphology, specifically the regulation of serially aligned sarcomeres, adapts more rapidly than parallel muscle morphology (i.e., muscle thickness, pennation angle) during immobilization and subsequent recovery in young and old rats. While SSN recovered slower in old rats compared to young, this recovery was still quicker than that observed for parallel muscle morphology, hence old rats retain a better ability to recover contractile tissue in series than in parallel. While this faster recovery of SSN did not rescue maximal force production, it did partially rescue the shape of the active torque-angle relationship. This rapid recovery of SSN represents critical early adaptive mechanisms driving longitudinal muscle growth to maintain functional capacity in older adults. Focusing on SSN growth during recovery from disuse or at baseline in older individuals therefore represents an appealing technique to help improve functional capacity.

## Acknowledgements

This project was supported by the Natural Sciences and Engineering Research Council of Canada (NSERC). The animals were obtained from the National Institute on Aging (NIA) aged rodent colonies.

## Conflict of interest statement

No conflicts of interest, financial or otherwise, are declared by the authors.

## Ethics statement

Approval was given by the University of Guelph’s Animal Care Committee and all protocols followed CCAC guidelines (AUP #4905).

## Data availability

All data generated or analysed during the study are available from the corresponding author upon request.

## Funding

This project was supported by the Natural Sciences and Engineering Research Council of Canada (NSERC), grant number RGPIN-2024-03782.

## Author contributions

A.H. and G.A.P. conceived and designed research; A.H. performed experiments; A.H. analyzed data; A.H. and G.A.P. interpreted results of experiments; A.H. prepared figures; A.H. and G.A.P. drafted manuscript; A.H. and G.A.P. edited and revised manuscript; A.H. and G.A.P. approved final version of manuscript.

---

**Supplemental Figure S1:**
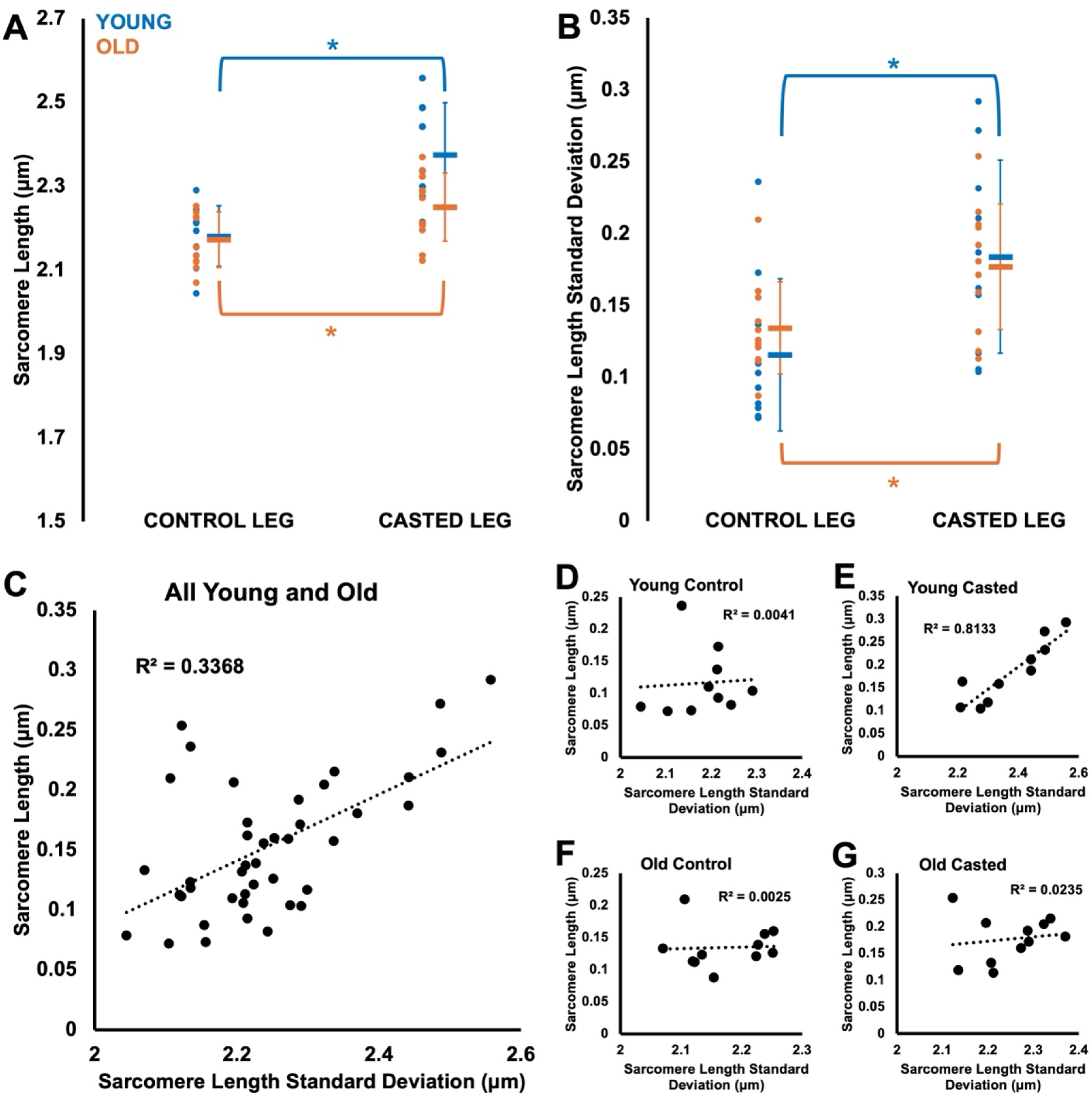
**A-B.** Differences in sarcomere length and sarcomere length standard deviation (estimate of sarcomere length non-uniformity) between control and casted legs in young (n = 10) and old (n = 11) rats, with post-cast and 4 wk recovery time points combined because there were effects of leg but not time. Data are displayed as mean ± standard deviation. *Difference between indicated points (*P* < 0.05). **C-G.** Relationships between sarcomere length and sarcomere length standard deviation. *Significant relationship (*P* < 0.05).

**Supplemental Figure S2:**
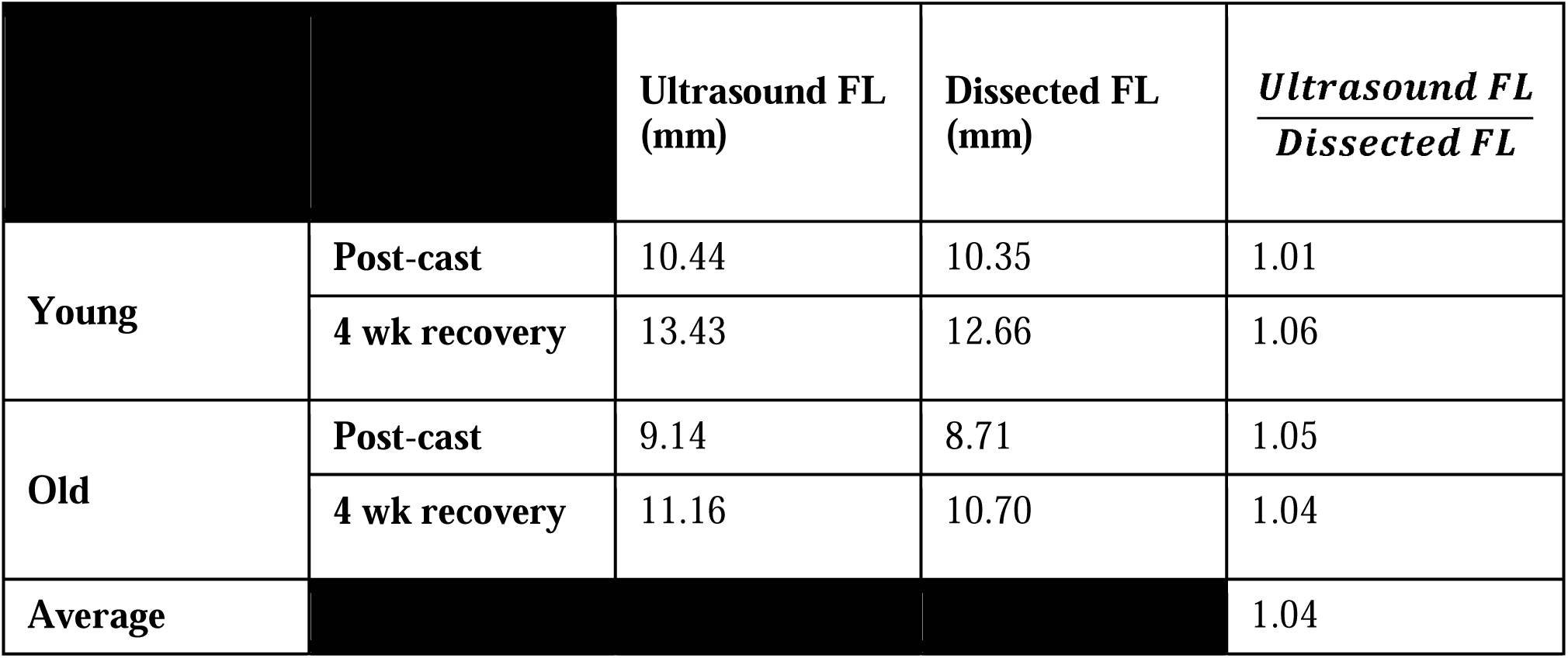
Ratio of fascicle length (FL) measured using ultrasound to FL measured on dissected fascicles from the same muscles

## Notes

### Competing Interest Statement

The authors have declared no competing interest.

## References

1. Lexell J, Henriksson-Larsén K, Winblad B, Sjöström M. Distribution of different fiber types in human skeletal muscles: Effects of aging studied in whole muscle cross sections. Muscle Nerve. 1983;6(8):588–95.

2. Frontera WR, Hughes VA, Fielding RA, Fiatarone MA, Evans WJ, Roubenoff R. Aging of skeletal muscle: a 12-yr longitudinal study. J Appl Physiol. 2000 Apr;88(4):1321–6.

3. Vandervoort AA. Aging of the human neuromuscular system. Muscle Nerve. 2002 Jan;25(1):17–25.

4. Delmonico MJ, Harris TB, Visser M, Park SW, Conroy MB, Velasquez-Mieyer P, et al. Longitudinal study of muscle strength, quality, and adipose tissue infiltration. Am J Clin Nutr. 2009 Dec 1;90(6):1579–85.

5. Power GA, Dalton BH, Rice CL. Human neuromuscular structure and function in old age: A brief review. J Sport Health Sci. 2013 Dec 1;2(4):215–26.

6. Narici MV, Maganaris CN, Reeves ND, Capodaglio P. Effect of aging on human muscle architecture. J Appl Physiol. 2003 Dec;95(6):2229–34.

7. Narici MV, Maffulli N. Sarcopenia: characteristics, mechanisms and functional significance. Br Med Bull. 2010 Sep 1;95(1):139–59.

8. Hinks A, Hawke TJ, Franchi MV, Power GA. The importance of serial sarcomere addition for muscle function and the impact of aging. J Appl Physiol [Internet]. 2023 Jul 6 [cited 2023 Jul 8]; Available from: https://journals.physiology.org/doi/abs/10.1152/japplphysiol.00205.2023

9. Hooper AC. Length, diameter and number of ageing skeletal muscle fibres. Gerontology. 1981;27(3):121–6.

10. Power GA, Crooks S, Fletcher JR, Macintosh BR, Herzog W. Age-related reductions in the number of serial sarcomeres contribute to shorter fascicle lengths but not elevated passive tension. J Exp Biol. 2021;224(10):jeb242172.

11. Hinks A, Patterson MA, Njai BS, Power GA. Age-related blunting of serial sarcomerogenesis and mechanical adaptations following 4 weeks of maximal eccentric resistance training. bioRxiv. 2023 Nov 10;

12. Butterfield TA, Herzog W. The magnitude of muscle strain does not influence serial sarcomere number adaptations following eccentric exercise. Pflüg Arch - Eur J Physiol. 2006 Feb;451(5):688–700.

13. Hinks A, Franchi MV, Power GA. The influence of longitudinal muscle fascicle growth on mechanical function. J Appl Physiol. 2022 Jul 1;133(1):87–103.

14. Hinks A, Jacob K, Mashouri P, Medak KD, Franchi MV, Wright DC, et al. Influence of weighted downhill running training on serial sarcomere number and work loop performance in the rat soleus. Biol Open. 2022 Jul 15;11(7):bio059491.

15. Narici MV, Maganaris CN. Adaptability of elderly human muscles and tendons to increased loading. J Anat. 2006 Apr;208(4):433–43.

16. Thom JM, Morse CI., Birch KM, Narici MV. Influence of muscle architecture on the torque and power–velocity characteristics of young and elderly men. Eur J Appl Physiol. 2007 Jun 26;100(5):613–9.

17. Narici M, Franchi M, Maganaris C. Muscle structural assembly and functional consequences. J Exp Biol. 2016 Jan;219(2):276–84.

18. Tabary JC, Tabary C, Tardieu C, Tardieu G, Goldspink G. Physiological and structural changes in the cat’s soleus muscle due to immobilization at different lengths by plaster casts. J Physiol. 1972 Jul 1;224(1):231–44.

19. Williams PE, Goldspink G. Changes in sarcomere length and physiological properties in immobilized muscle. J Anat. 1978;127(3):459–68.

20. Witzmann FA, Kim DH, Fitts RH. Hindlimb immobilization: length-tension and contractile properties of skeletal muscle. J Appl Physiol. 1982 Aug 1;53(2):335–45.

21. Heslinga JW, Huijing PA. Muscle length-force characteristics in relation to muscle architecture: a bilateral study of gastrocnemius medialis muscles of unilaterally immobilized rats. Eur J Appl Physiol. 1993 Apr;66(4):289–98.

22. Surkan MJ, Gibson W. Interventions to Mobilize Elderly Patients and Reduce Length of Hospital Stay. Can J Cardiol. 2018 Jul;34(7):881–8.

23. Valenzuela PL, Morales JS, Pareja-Galeano H, Izquierdo M, Emanuele E, de la Villa P, et al. Physical strategies to prevent disuse-induced functional decline in the elderly. Ageing Res Rev. 2018 Nov;47:80–8.

24. Williams PE, Goldspink G. The effect of immobilization on the longitudinal growth of striated muscle fibres. J Anat. 1973;116(1):45–55.

25. Koh TJ, Tidball JG. Nitric oxide synthase inhibitors reduce sarcomere addition in rat skeletal muscle. J Physiol. 1999 Aug;519(1):189–96.

26. Dayanidhi S, Kinney MC, Dykstra PB, Lieber RL. Does a Reduced Number of Muscle Stem Cells Impair the Addition of Sarcomeres and Recovery from a Skeletal Muscle Contracture? A Transgenic Mouse Model. Clin Orthop. 2020 Apr;478(4):886–99.

27. Horner AM, Azizi E, Roberts TJ. The interaction of in vivo muscle operating lengths and passive stiffness in rat hindlimbs. J Exp Biol. 2024 Feb 14;jeb.246280.

28. Johannsen DL, DeLany JP, Frisard MI, Welsch MA, Rowley CK, Fang X, et al. Physical activity in aging: Comparison among young, aged, and nonagenarian individuals. J Appl Physiol. 2008 Aug;105(2):495–501.

29. Sun F, Norman IJ, While AE. Physical activity in older people: a systematic review. BMC Public Health. 2013 May 6;13(1):449.

30. Gomes M, Figueiredo D, Teixeira L, Poveda V, Paúl C, Santos-Silva A, et al. Physical inactivity among older adults across Europe based on the SHARE database. Age Ageing. 2017 Jan 19;46(1):71–7.

31. Hagen JL, Krause DJ, Baker DJ, Fu MH, Tarnopolsky MA, Hepple RT. Skeletal Muscle Aging in F344BN F1-Hybrid Rats: I. Mitochondrial Dysfunction Contributes to the Age-Associated Reduction in VO2max. J Gerontol Ser A. 2004 Nov 1;59(11):1099–110.

32. Suetta C, Hvid LG, Justesen L, Christensen U, Neergaard K, Simonsen L, et al. Effects of aging on human skeletal muscle after immobilization and retraining. J Appl Physiol Bethesda Md 1985. 2009 Oct;107(4):1172–80.

33. Suetta C, Frandsen U, Mackey AL, Jensen L, Hvid LG, Bayer ML, et al. Ageing is associated with diminished muscle re-growth and myogenic precursor cell expansion early after immobility-induced atrophy in human skeletal muscle. J Physiol. 2013 Aug 1;591(15):3789–804.

34. Zarzhevsky N, Menashe O, Carmeli E, Stein H, Reznick AZ. Capacity for recovery and possible mechanisms in immobilization atrophy of young and old animals. Ann N Y Acad Sci. 2001 Apr;928:212–25.

35. de la Tour EH, Tabary JC, Tabary C, Tardieu C. The respective roles of muscle length and muscle tension in sarcomere number adaptation of guinea-pig soleus muscle. J Physiol (Paris). 1979;75(5):589–92.

36. Kinney MC, Dayanidhi S, Dykstra PB, McCarthy JJ, Peterson CA, Lieber RL. Reduced skeletal muscle satellite cell number alters muscle morphology after chronic stretch but allows limited serial sarcomere addition: Satellite Cells and Sarcomere Addition. Muscle Nerve. 2017 Mar;55(3):384–92.

37. Hinks A, Franchi MV, Power GA. Ultrasonographic measurements of fascicle length overestimate adaptations in serial sarcomere number. Exp Physiol. 2023 Aug 23;108(10):1308–24.

38. Aoki MS, Soares AG, Miyabara EH, Baptista IL, Moriscot AS. Expression of genes related to myostatin signaling during rat skeletal muscle longitudinal growth: Myostatin and Longitudinal Growth. Muscle Nerve. 2009 Dec;40(6):992–9.

39. Warren GL, Stallone JL, Allen MR, Bloomfield SA. Functional recovery of the plantarflexor muscle group after hindlimb unloading in the rat. Eur J Appl Physiol. 2004 Oct;93(1– 2):130–8.

40. Padilla CJ, Harrigan ME, Harris H, Schwab JM, Rutkove SB, Rich MM, et al. Profiling age-related muscle weakness and wasting: neuromuscular junction transmission as a driver of age-related physical decline. GeroScience. 2021 Jun 1;43(3):1265–81.

41. Chen J, Mashouri P, Fontyn S, Valvano M, Elliott-Mohamed S, Noonan AM, et al. The influence of training-induced sarcomerogenesis on the history dependence of force. J Exp Biol. 2020 Jun 19;223(Pt 15):jeb218776.

42. Mele A, Fonzino A, Rana F, Camerino GM, De Bellis M, Conte E, et al. In vivo longitudinal study of rodent skeletal muscle atrophy using ultrasonography. Sci Rep. 2016 Apr;6(1):20061.

43. Franchi MV, Fitze DP, Raiteri BJ, Hahn D, Spörri J. Ultrasound-derived Biceps Femoris Long-Head Fascicle Length: Extrapolation Pitfalls. Med Sci Sports Exerc. 2019 Aug;52(1):233–43.

44. Peixinho CC, Ribeiro MB, Resende CMC, Werneck-de-Castro JPS, de Oliveira LF, Machado JC. Ultrasound biomicroscopy for biomechanical characterization of healthy and injured triceps surae of rats. J Exp Biol. 2011 Nov 15;214(22):3880–6.

45. Akagi R, Hinks A, Power GA. Differential changes in muscle architecture and neuromuscular fatigability induced by isometric resistance training at short and long muscle-tendon unit lengths. J Appl Physiol. 2020 Jun 18;129(1):173–84.

46. Butterfield TA, Leonard TR, Herzog W. Differential serial sarcomere number adaptations in knee extensor muscles of rats is contraction type dependent. J Appl Physiol. 2005;99:7.

47. Lieber RL, Yeh Y, Baskin RJ. Sarcomere length determination using laser diffraction. Effect of beam and fiber diameter. Biophys J. 1984 May;45(5):1007–16.

48. Leeuwenburgh C, Gurley CM, Strotman BA, Dupont-Versteegden EE. Age-related differences in apoptosis with disuse atrophy in soleus muscle. Am J Physiol Regul Integr Comp Physiol. 2005 May;288(5):R1288–1296.

49. Fuqua JD, Lawrence MM, Hettinger ZR, Borowik AK, Brecheen PL, Szczygiel MM, et al. Impaired proteostatic mechanisms other than decreased protein synthesis limit old skeletal muscle recovery after disuse atrophy. J Cachexia Sarcopenia Muscle. 2023 Jul 14;14(5):2076–89.

50. Hepple RT, Rice CL. Innervation and neuromuscular control in ageing skeletal muscle. J Physiol. 2016 Apr 15;594(8):1965–78.

51. Sakuma K, Aoi W, Yamaguchi A. Current understanding of sarcopenia: possible candidates modulating muscle mass. Pflüg Arch - Eur J Physiol. 2015 Feb 1;467(2):213–29.

52. Larsson L, Degens H, Li M, Salviati L, Lee YI, Thompson W, et al. Sarcopenia: Aging-Related Loss of Muscle Mass and Function. Physiol Rev. 2019 Jan 1;99(1):427–511.

53. Kubo K, Kanehisa H, Azuma K, Ishizu M, Kuno SY, Okada M, et al. Muscle Architectural Characteristics in Young and Elderly Men and Women. Int J Sports Med. 2003 Feb;24(2):125–30.

54. Morse CI, Thom JM, Birch KM, Narici MV. Changes in triceps surae muscle architecture with sarcopenia. Acta Physiol Scand. 2005 Mar;183(3):291–8.

55. Baudry S, Lecoeuvre G, Duchateau J. Age-related changes in the behavior of the muscle-tendon unit of the gastrocnemius medialis during upright stance. J Appl Physiol Bethesda Md 1985. 2012 Jan;112(2):296–304.

56. Stenroth L, Peltonen J, Cronin NJ, Sipilä S, Finni T. Age-related differences in Achilles tendon properties and triceps surae muscle architecture in vivo. J Appl Physiol Bethesda Md 1985. 2012 Nov;113(10):1537–44.

57. Power GA, Makrakos DP, Rice CL, Vandervoort AA. Enhanced force production in old age is not a far stretch: an investigation of residual force enhancement and muscle architecture. Physiol Rep. 2013 Jun;1(1):e00004.

58. Panizzolo FA, Green DJ, Lloyd DG, Maiorana AJ, Rubenson J. Soleus fascicle length changes are conserved between young and old adults at their preferred walking speed. Gait Posture. 2013 Sep;38(4):764–9.

59. Baroni BM, Geremia JM, Rodrigues R, Borges MK, Jinha A, Herzog W, et al. Functional and Morphological Adaptations to Aging in Knee Extensor Muscles of Physically Active Men. J Appl Biomech. 2013 Oct 1;29(5):535–42.

60. Wu R, Delahunt E, Ditroilo M, Lowery M, De Vito G. Effects of age and sex on neuromuscular-mechanical determinants of muscle strength. Age Dordr Neth. 2016 Jun;38(3):57.

61. Conway KA, Franz JR. Shorter gastrocnemius fascicle lengths in older adults associate with worse capacity to enhance push-off intensity in walking. Gait Posture. 2020 Mar;77:89–94.

62. Kelp NY, Gore A, Clemente CJ, Tucker K, Hug F, Dick TJM. Muscle architecture and shape changes in the gastrocnemii of active younger and older adults. J Biomech. 2021 Dec 2;129:110823.

63. Karamanidis K, Arampatzis A. Mechanical and morphological properties of different muscle–tendon units in the lower extremity and running mechanics: effect of aging and physical activity. J Exp Biol. 2005 Oct 15;208(20):3907–23.

64. Barber LA, Barrett RS, Gillett JG, Cresswell AG, Lichtwark GA. Neuromechanical properties of the triceps surae in young and older adults. Exp Gerontol. 2013 Nov;48(11):1147–55.

65. Erskine RM, Tomlinson DJ, Morse CI, Winwood K, Hampson P, Lord JM, et al. The individual and combined effects of obesity-and ageing-induced systemic inflammation on human skeletal muscle properties. Int J Obes 2005. 2017 Jan;41(1):102–11.

66. Gerstner GR, Thompson BJ, Rosenberg JG, Sobolewski EJ, Scharville MJ, Ryan ED. Neural and Muscular Contributions to the Age-Related Reductions in Rapid Strength. Med Sci Sports Exerc. 2017 Jul;49(7):1331–9.

67. Franchi MV, Monti E, Carter A, Quinlan JI, Herrod PJJ, Reeves ND, et al. Bouncing Back! Counteracting Muscle Aging With Plyometric Muscle Loading. Front Physiol. 2019 Mar 5;10:178.

68. Quinlan JI, Franchi MV, Gharahdaghi N, Badiali F, Francis S, Hale A, et al. Muscle and tendon adaptations to moderate load eccentric vs. concentric resistance exercise in young and older males. GeroScience. 2021;43(4):1567–84.

69. Pinel S, Kelp NY, Bugeja JM, Bolsterlee B, Hug F, Dick TJM. Quantity versus quality: Age-related differences in muscle volume, intramuscular fat, and mechanical properties in the triceps surae. Exp Gerontol. 2021 Dec;156:111594.

70. Power GA, Crooks S, Fletcher JR, Macintosh BR, Herzog W. Age-related reductions in the number of serial sarcomeres contribute to shorter fascicle lengths but not elevated passive tension. 2020 Dec 22 [cited 2021 May 18]; Available from: http://biorxiv.org/lookup/doi/10.1101/2020.12.21.423814

71. Ansved T. Effects of immobilization on the rat soleus muscle in relation to age. Acta Physiol Scand. 1995 Jul;154(3):291–302.

72. Fisher JS, Brown M. Immobilization effects on contractile properties of aging rat skeletal muscle. Aging Milan Italy. 1998 Feb;10(1):59–66.

73. Hvid L, Aagaard P, Justesen L, Bayer ML, Andersen JL, Ørtenblad N, et al. Effects of aging on muscle mechanical function and muscle fiber morphology during short-term immobilization and subsequent retraining. J Appl Physiol Bethesda Md 1985. 2010 Dec;109(6):1628–34.

74. Hvid LG, Suetta C, Nielsen JH, Jensen MM, Frandsen U, Ørtenblad N, et al. Aging impairs the recovery in mechanical muscle function following 4 days of disuse. Exp Gerontol. 2014 Apr;52:1–8.

75. Herbert RD, Gandevia SC. The passive mechanical properties of muscle. J Appl Physiol. 2019 May 1;126(5):1442–4.

76. Binder-Markey BI, Sychowski D, Lieber RL. Systematic review of skeletal muscle passive mechanics experimental methodology. J Biomech. 2021 Dec;129:110839.

77. Palokangas H, Kovanen V, Duncan A, Robins SP. Age-related changes in the concentration of hydroxypyridinium crosslinks in functionally different skeletal muscles. Matrix Stuttg Ger. 1992 Aug;12(4):291–6.

78. Gosselin LE, Martinez DA, Vailas AC, Sieck GC. Passive length-force properties of senescent diaphragm: relationship with collagen characteristics. J Appl Physiol Bethesda Md 1985. 1994 Jun;76(6):2680–5.

79. Haus JM, Carrithers JA, Trappe SW, Trappe TA. Collagen, cross-linking, and advanced glycation end products in aging human skeletal muscle. J Appl Physiol Bethesda Md 1985. 2007 Dec;103(6):2068–76.

80. Olson LC, Nguyen TM, Heise RL, Boyan BD, Schwartz Z, McClure MJ. Advanced Glycation End Products Are Retained in Decellularized Muscle Matrix Derived from Aged Skeletal Muscle. Int J Mol Sci. 2021 Aug 17;22(16):8832.

81. Brashear SE, Wohlgemuth RP, Hu LY, Jbeily EH, Christiansen BA, Smith LR. Collagen cross-links scale with passive stiffness in dystrophic mouse muscles, but are not altered with administration of a lysyl oxidase inhibitor. PloS One. 2022;17(10):e0271776.

82. Wisdom KM, Delp SL, Kuhl E. Use it or lose it: multiscale skeletal muscle adaptation to mechanical stimuli. Biomech Model Mechanobiol. 2015 Apr;14(2):195–215.

83. Koh TJ, Herzog W. Excursion is important in regulating sarcomere number in the growing rabbit tibialis anterior. J Physiol. 1998 Apr 1;508 (Pt 1)(Pt 1):267–80.

84. Soares AG, Aoki MS, Miyabara EH, DeLuca CV, Ono HY, Gomes MD, et al. Ubiquitin-ligase and deubiquitinating gene expression in stretched rat skeletal muscle. Muscle Nerve. 2007 Nov;36(5):685–93.

85. Seo DY, Hwang BG. Effects of exercise training on the biochemical pathways associated with sarcopenia. Phys Act Nutr. 2020 Sep 30;24(3):32–8.

86. González-Blanco L, Bermúdez M, Bermejo-Millo JC, Gutiérrez-Rodríguez J, Solano JJ, Antuña E, et al. Cell interactome in sarcopenia during aging. J Cachexia Sarcopenia Muscle. 2022;13(2):919–31.

87. Gallegly JC, Turesky NA, Strotman BA, Gurley CM, Peterson CA, Dupont-Versteegden EE. Satellite cell regulation of muscle mass is altered at old age. J Appl Physiol Bethesda Md 1985. 2004 Sep;97(3):1082–90.

88. Snijders T, Nederveen JP, McKay BR, Joanisse S, Verdijk LB, van Loon LJC, et al. Satellite cells in human skeletal muscle plasticity. Front Physiol. 2015;6:283.

89. Huo F, Liu Q, Liu H. Contribution of muscle satellite cells to sarcopenia. Front Physiol. 2022 Aug 12;13:892749.

90. Efimenko A, Starostina E, Kalinina N, Stolzing A. Angiogenic properties of aged adipose derived mesenchymal stem cells after hypoxic conditioning. J Transl Med. 2011 Jan 18;9:10.

91. Hettinger ZR, Hamagata K, Confides AL, Lawrence MM, Miller BF, Butterfield TA, et al. Age-Related Susceptibility to Muscle Damage Following Mechanotherapy in Rats Recovering From Disuse Atrophy. J Gerontol A Biol Sci Med Sci. 2021 Jun 28;76(12):2132–40.

92. Brooks SV, Faulkner JA. Contraction-induced injury: recovery of skeletal muscles in young and old mice. Am J Physiol. 1990 Mar;258(3 Pt 1):C436–442.

93. Nunes EA, Stokes T, McKendry J, Currier BS, Phillips SM. Disuse-induced skeletal muscle atrophy in disease and nondisease states in humans: mechanisms, prevention, and recovery strategies. Am J Physiol-Cell Physiol. 2022 Jun;322(6):C1068–84.

